# The crystal structure of vaccinia virus protein E2 and perspectives on the prediction of novel viral protein folds

**DOI:** 10.1101/2021.10.14.464338

**Authors:** William N.D. Gao, Chen Gao, Janet E. Deane, David C.J. Carpentier, Geoffrey L. Smith, Stephen C. Graham

## Abstract

The morphogenesis of vaccinia virus (VACV, family *Poxviridae*), the smallpox vaccine, is a complex process involving multiple distinct cellular membranes and resulting in multiple different forms of infectious virion. Efficient release of enveloped virions, which promote systemic spread of infection within hosts, requires the VACV protein E2 but the molecular basis of E2 function remains unclear and E2 lacks sequence homology to any well-characterised family of proteins. We solved the crystal structure of VACV E2 to 2.3 Å resolution, revealing that it comprises two domains with novel folds: an N-terminal annular (ring) domain and a C-terminal head domain. The C-terminal head domain displays weak structural homology with cellular (pseudo)kinases but lacks conserved surface residues or kinase features, suggesting that it is not enzymatically active, and possesses a large surface basic patch that might interact with phosphoinositide lipid headgroups. Recent deep learning methods have revolutionised our ability to predict the three-dimensional structures of proteins from primary sequence alone. VACV E2 is an exemplar ‘difficult’ viral protein target for structure prediction, being comprised of multiple novel domains and lacking sequence homologues outside *Poxviridae*. AlphaFold2 nonetheless succeeds in predicting the structures of the head and ring domains with high and moderate accuracy, respectively, allowing accurate inference of multiple structural properties. The advent of highly accurate virus structure prediction marks a step-change in structural virology and beckons a new era of structurally-informed molecular virology.

---

Vaccinia virus (VACV) is the prototype member of the *Poxviridae*, a family of DNA viruses producing large and complex enveloped virions [1]. The family includes variola virus, the causative agent of the highly infectious and lethal disease smallpox, and several viruses endemic in a variety of animal species, some linked with increasing incidences of zoonotic spread and disease in humans [2–4]. While a concerted vaccination programme led to the WHO declaring smallpox eradicated in 1980, the potential for re-emergence of poxvirus disease remains and only two drugs, TPOXX and Tembexa, are licenced for the treatment of orthopoxvirus infection.

Orthopoxviruses produce of two distinct types of infectious virion, mature virions (MVs, also called intracellular mature virions, IMVs) and enveloped virions (EVs, also known as extracellular enveloped virions EEVs). MVs are formed in cytoplasmic viral factories, where the genome-containing viral core and lateral bodies are wrapped by a single lipid membrane derived from the endoplasmic reticulum [5]. MVs are highly stable and, when released upon cell lysis, can survive in the environment to mediate horizontal spread to new hosts. However, MVs are susceptible to recognition by host adaptive immune response due to the abundance of conserved viral epitopes on their surface, including components of the virus membrane fusion and entry machinery. Prior to cell lysis a proportion of MVs are trafficked on microtubules to sites enriched in trans-Golgi/early endosome derived membranes, where they are wrapped by two additional envelopes to form intracellular enveloped virus (IEV, also known as wrapped virus, WV). These IEVs recruit the cellular kinesin-1 microtubule-associated motor complex to mediate virion trafficking to the cell periphery [6–9], whereupon the outer IEV envelope fuses with the cell membrane to release EVs onto the cell surface and into the extracellular medium. These EVs play an important role in cell-to-cell and systemic spread of infection within a host [10].

During IEV egress at least three viral proteins are involved in the activation of kinesin-1-dependent transport of IEVs. These include an integral membrane protein A36 and two cytoplasmic proteins, F12 and E2. Kinesin-1 is a tetramer of two heavy chains, comprising the microtubule-binding motor domain and a long coiled-coil dimerization domain, and two light chains, each comprising a tetratricopeptide repeat (TPR) domain and a coiled-coil domain that mediates dimerization plus heavy-chain association. A36 is associated with the outer IEV envelope [11] and possesses two tryptophan acidic (WE/WD) motifs, conserved in cellular kinesin light chain (KLC) binding proteins [12], that associate with a binding groove in the KLC TPR domain [13]. E2 also associates with KLC, binding to the unstructured C-terminal tail present on a subset of KLC isoforms [14, 15]. E2 and F12 function as a complex and both are essential for IEV egress [16]. The E2:F12 complex may associate with IEVs through an interaction between A36 and F12 [17], though the maintenance of E2:F12-mediated IEV egress in the absence of A36 [18] suggests that E2:F12 may utilise additional/alternative interactions to bind IEVs. The molecular basis by which E2 and F12 regulate the recruitment of kinesin-1 and promote microtubule-based IEV trafficking remains poorly understood.

While conserved across poxviruses, VACV E2 lacks identifiable sequence homology to any other protein family. Viral proteins in general, and poxvirus proteins in particular, can maintain structural homology to proteins of known function in the absence of identifiable sequence similarity [19–21]. We therefore sought to solve the structure of E2 from VACV strain Western Reserve. As extensive attempts to express E2 in bacterial (*Escherichia coli*) and insect cell systems were largely unsuccessful, a mammalian expression system was pursued. Small-scale expression tests using transient transfection of codon optimised E2 into human embryonic kidney (HEK)293T cells confirmed successful purification using cobalt affinity chromatography of VACV E2 tagged at the carboxy terminus with a decahistidine tag. Large-scale expression was thus performed via transient transfection of Freestyle 293F cells cultured in suspension and purification via cobalt affinity chromatography and size exclusion chromatography (SEC) followed by anion exchange chromatography, yielding ^~^0.75 mg of highly pure E2 per L of cultured cells (Figure 1A). Differential scanning fluorimetry (a.k.a. Thermofluor) confirmed that the protein was folded (Figure 1B), the biphasic melt curve of E2 suggesting the presence of two independently-folded domains. SEC analysis with inline multiangle light scattering (SEC-MALS) confirmed that the protein is predominantly monomeric (Figure 1C), the observed molecular mass (92.3 kDa) being close to expected mass as calculated from the sequence (87.5 kDa).

**Figure 1.**
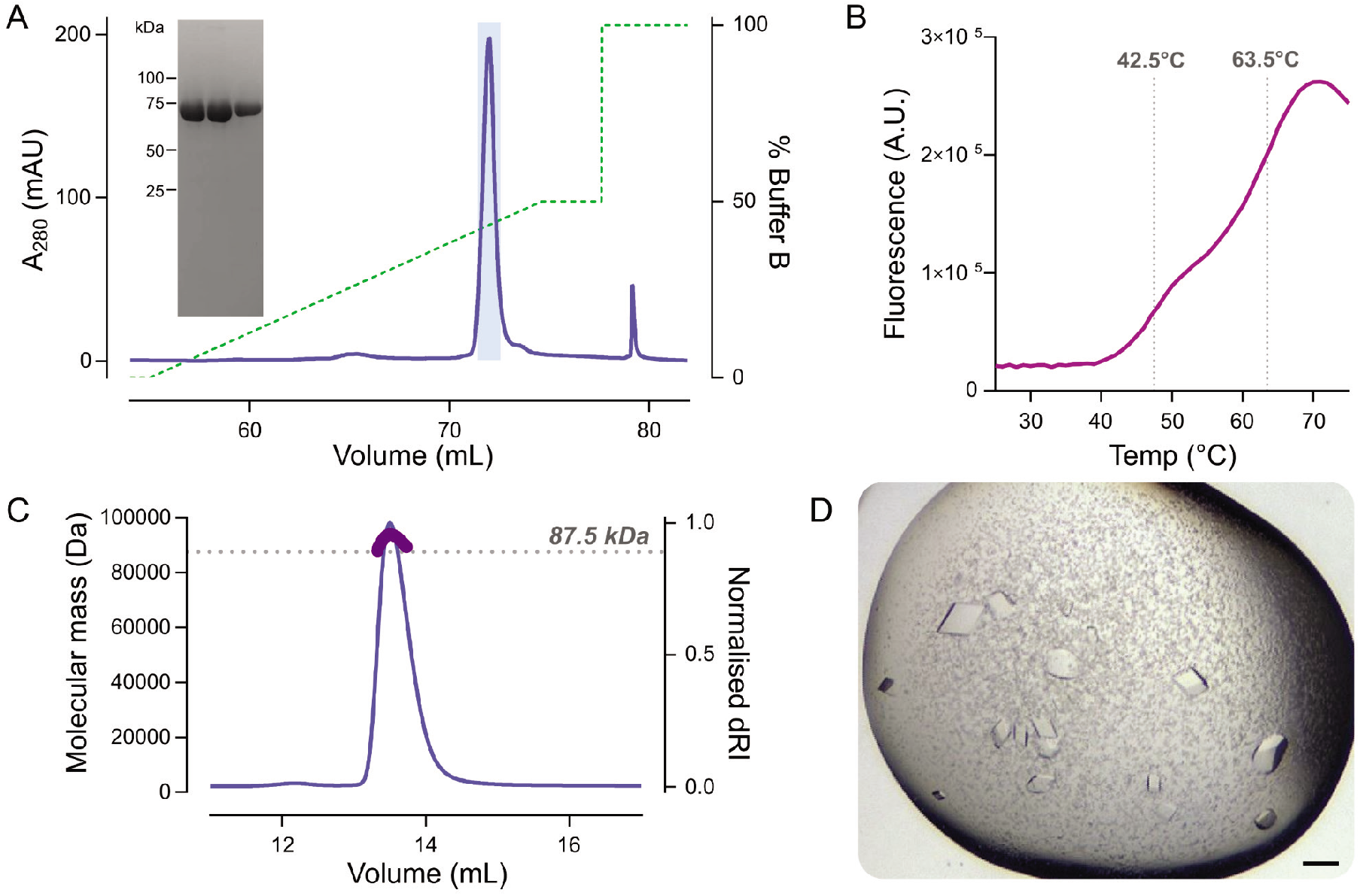
Purification, characterisation and crystallisation of VACV E2. (A) Preparative anion exchange chromatography. VACV E2 was expressed in Freestyle 293F cells and grown in Freestyle 293 medium (ThermoFisher) as per the manufacturer’s instructions, by transfection of pcDNA3 encoding VACV E2 with a C-terminal A_3_H_10_ tag mixed in a 1:2 ratio with 25 kDa branched polyethylenimine (PEI), adding 1 μg DNA and 2 μg PEI per mL of cultured cells. Cells were cultured for 40 h in a humidified 8% CO_2_ atmosphere at 37°C before being harvested by centrifugation, washed thrice with ice-cold PBS, resuspended in lysis buffer (50 mM Tris pH 8.0, 150 mM NaCl supplemented with protease inhibitors [Roche]) and lysed by five passages through a 23G needle. Lysates were clarified by centrifugation (40,000×g, 40 min, 4°C) before being applied to a 5 mL HiTrap TALON Crude Co^2+^ affinity column (Cytiva) and purified with elution in 200 mM imidazole as per the manufacturer’s instructions. Pooled eluate was further purified by size-exclusion chromatography (SEC) using a Superdex 200 10/300 GL column (Cytiva) equilibrated in SEC buffer (20 mM Tris pH 8.0, 200 mM NaCl, 1 mM DTT). As eluted protein retained contaminants, E2 was further purified by anion exchange chromatography with a MonoQ 5/50 GL column (Cytiva) using a linear gradient of 0–500 mM NaCl (green dashed line) in 20 mM Tris pH 8.0, protein elution being monitored using UV absorbance (blue line). Peak fractions containing VACV E2 that were pooled and used for subsequent analysis are highlighted (light blue) and SDS-PAGE of these fractions shows VACV E2 to be highly pure. (B) Differential scanning fluorimetry of VACV E2. Purified E2 (4 μg) was mixed with 1× Protein Thermal Shift dye (Applied Biosystems) in a final volume of 20 μL and heated from 25 to 95°C at 1 degree per 30 s, with fluorescence (purple curve) being monitored at each increment. Two inflection points are visible (grey dotted lines), consistent with biphasic melting. (C) SEC with inline multi-angle light scattering (SEC-MALS) shows VACV E2 to be predominantly monomeric. Purified E2 (100 μg) was injected onto a Superdex 200 10/300 GL column (Cytiva) equilibrated in SEC buffer at 0.4 mL/min at room temperature with inline measurement of static light scattering (DAWN 8+, Wyatt Technology), differential refractive index (dRI; Optilab T-rEX, Wyatt Technology), and 280 nm absorbance (Agilent 1260 UV, Agilent Technologies). The normalised dRI is shown (thin blue line), as is the molecular mass of the peak (thick purple line) as calculated using ASTRA6 (Wyatt Technology) assuming a protein dn/dc of 0.186. The calculated mass (92.3 kDa) is in good agreement with theoretical mass of VACV E2 with a C-terminal A_3_H_10_ tag (87.5 kDa; dotted grey line), confirming that the protein is predominantly monomeric. (D) Crystals of VACV E2, grown by sitting drop vapour diffusion. 200 nL of 9.7 mg/mL E2 was mixed with 60 nL of reservoir solution (50 mM ADA (N-(2-acetamido)iminodiacetic acid) pH 6.5, 50 mM ADA pH 7.0, 10% v/v 2-methyl-2,4-pentanediol [MPD]) and equilibrated against 80 μL reservoirs at 20°C, crystals growing within 21 days. Scale bar = 100 μm.

VACV E2 at a concentration of 10.85 mg/mL was subjected to nanolitre crystallisation trials, crystals being obtained when 200 nL of protein was mixed with an equal volume of reservoir solution and equilibrated against an 80 μL reservoir of 0.1 M sodium formate pH 7.0, 12% (w/v) polyethylene glycol 3350 at 20°C. Crystals were cryoprotected by brief immersion in reservoir solution supplemented with 25% (v/v) glycerol before plunge cryocooling and diffraction data were recorded at Diamond Light Source beamline I04-1 to 2.7 Å resolution. Unfortunately, all crystals obtained in the first crystallisation experiment were consumed in the process of obtaining this diffraction dataset and the extensive attempts to reproduce crystallisation under these conditions were unsuccessful, precluding solution of the phase problem by experimental methods. Attempts to solve the structure of E2 via molecular replacement using *ab initio* models generated by the I-TASSER modelling server [22], which was state-of-the-art at the time (in 2016), were unsuccessful. Exhaustive molecular replacement searches using 68,087 folded domains drawn from across the PDB [23] were also unsuccessful, suggesting that E2 either possessed a novel fold or that the structural homology to the (multiple) domains was insufficient to facilitate structure solution.

Extensive sparse matrix screening eventually identified new conditions for the crystallisation of VACV E2 (Figure 1D), these new crystals sharing the same space group and unit cell dimensions as the crystal that was collected previously. The structure of VACV E2 was solved by single isomorphous replacement using an ethylmercurithiosalicylate (EMTS, a.k.a. Thimerosal) derivative and the structure was refined to 2.3 Å resolution (Table 1). The structure of VACV E2 comprises two folded domains: an N-terminal annular (ring) domain spanning residues 1–454 and C-terminal compact globular (head) domain spanning residues 455–732 (Figure 2A). The final 5 amino acids of E2 (FKSSK) and all residues of the cloning tag were not visible in electron density and are presumed disordered. E2 is primarily α-helical, with only three short β-sheets being evident at the apex of the head domain (Figure 2B). Structural homology searches performed using DALI [24] and PDBePISA [25] did not identify any significant structural homologues of the ring domain. The head domain does share weak homology to protein and glycan kinase domains, and to the bacterial SidJ pseudokinase domain that possesses glutamylase activity [26]. However, the overall structural correspondence is low, with less than half of the domain structurally aligned, and key kinase catalytic motifs such as the glycine-rich loop and catalytic lysine and aspartic acid residues are not conserved. Mapping the conservation of E2 sequence across poxviruses onto the structure [27] does not reveal any surface patches of high conservation as would be expected at an enzyme active site. This suggests that the E2 head domain lacks catalytic activity and that any similarity between this head domain and (pseudo)kinases is either spurious and/or vestigial, potentially representing a cellular (pseudo)kinase domain acquired by an ancestral poxvirus that has subsequently evolved toward a novel function [20, 21].

**Table 1.** X-ray diffraction data collection and structure refinement. Crystals of VACV E2 were grown by sitting drop vapour diffusion against 80 μL reservoirs, crystallisation drops containing 200 nL 9.7 mg/mL E2 plus 120 nL 50 mM ADA pH 6.0, 50 mM ADA pH 6.5, 8% v/v MPD (*Low-resolution native*), 120 nL 50 mM ADA pH 6.0, 50 mM ADA pH 6.5, 8% v/v MPD (*EMTS soak*), or 60 nL 50 mM ADA pH 6.5, 50 mM ADA pH 7.0, 10% v/v 2-methyl-2,4-pentanediol (*High-resolution native*). Heavy atom derivitisation was achieved by soaking crystals for 90 min in reservoir solution supplemented with 1 mM ethylmercurithiosalicylate (EMTS) and 25% glycerol. All crystals were cryoprotected by rapid transfer to reservoir solution supplemented with 25% v/v glycerol before plunge cryocooling in liquid nitrogen. Diffraction data were recorded at Diamond Light Source beamline I04 and processed using DIALS [44] as implemented in the xia2 [45] autoprocessing pipeline. The structure of VACV E2 was solved via single isomorphous replacement with anomalous scattering (SIRAS) by CRANK2 [46] using the *low-resolution native* and *EMTS soak* datasets. The initial model was then used to phase the *high-resolution native* data and the model was completed and refined using COOT [47], autoBUSTER [48] and phenix.refine [49] in consultation with MolProbity [50] and the validation tools present in COOT [47]. Values in parentheses refer to the high-resolution shell. The atomic coordinates and structure factors have been deposited in the Protein Data Bank [51] with accession code 7PHY and the original diffraction data are available from the University of Cambridge Apollo repository (https://doi.org/10.17863/CAM.74391).

| Data collection | *Low resolution native* | *EMTS soak* | *High-resolution native* |
| --- | --- | --- | --- |
| Wavelength (Å) | 0.9795 | 0.9795 | 0.9795 |
| Space group | *P* 2<sub>1</sub>2<sub>1</sub>2<sub>1</sub> | *P* 2<sub>1</sub>2<sub>1</sub>2<sub>1</sub> | *P* 2<sub>1</sub>2<sub>1</sub>2<sub>1</sub> |
| Cell dimensions (*a*, *b*, *c*) (Å) |  | 78.08, 90.52, 144.23 | 77.17, 90.93, 147.20 |
| Resolution range (Å) | 146.3–2.5<br>(2.51–2.47) | 59.1–3.1<br>(3.11–3.06) | 39.2–2.3<br>(2.34–2.30) |
| Completeness (%) | 100.0 (99.8) | 100.0 (98.6) | 100.0 (100.0) |
| Multiplicity | 13.0 (13.4) | 12.8 (12.9) | 13.2 (12.4) |
| CC<sub>1/2</sub> | 0.997 (0.618) | 0.999 (0.509) | 1.00 (0.404) |
| Mean I/$\sigma$(I) | 15.98 (2.48) | 8.6 (2.3) | 19.0 (0.6) |
| *R*<sub>merge</sub> | 0.098 (0.893) | 0.214 (0.937) | 0.074 (2.422) |
| *R*<sub>meas</sub> | 0.102 (0.928) | 0.223 (0.976) | 0.077 (2.526) |
| *R*<sub>pim</sub> | 0.028 (0.251) | 0.062 (0.269) | 0.021 (0.708) |
| Anomalous completeness (%) | 100.0 (99.8) | 99.9 (98.4) | 100.0 (99.7) |
| Anomalous multiplicity | 6.9 (6.9) | 6.8 (6.8) | 6.9 (6.5) |
| Anomalous CC<sub>1/2</sub> | -0.019 (-0.015) | 0.280 (-0.010) | -0.136 (0.021) |
| Wilson B factor (Å<sup>2</sup>) | 52.0 | 68.2 | 58.9 |
| **Refinement** |  |  |  |
| Resolution range (Å) |  |  | 35.5–2.3<br>(2.35–2.30) |
| Reflections |  |  |  |
| Working set |  |  | 44,190 (2543) |
| Test set |  |  | 2396 (153) |
| *R* |  |  | 0.1943 (0.4000) |
| *R*<sub>free</sub> |  |  | 0.2370 (0.4195) |
| No. of atoms |  |  |  |
| Protein |  |  | 6005 |
| Solvent |  |  | 249 |
| Other<sup>a</sup> |  |  | 41 |
| Root mean square deviation |  |  |  |
| Bond length (Å) |  |  | 0.008 |
| Bond angle (°) |  |  | 0.866 |
| Ramachandran favoured (%) |  |  | 97.81 |
| Ramachandran outliers (%) |  |  | 0.14 |
| Clash score |  |  | 2.46 |
| Poor rotamers (%) |  |  | 0.58 |
| Mean B value (Å<sup>2</sup>) |  |  | 75.71 |

<sup>a</sup>The N terminus of E2 was modelled as an N-acetylated methionine and five ordered glycerol molecules were observed in the structure.

**Figure 2.**
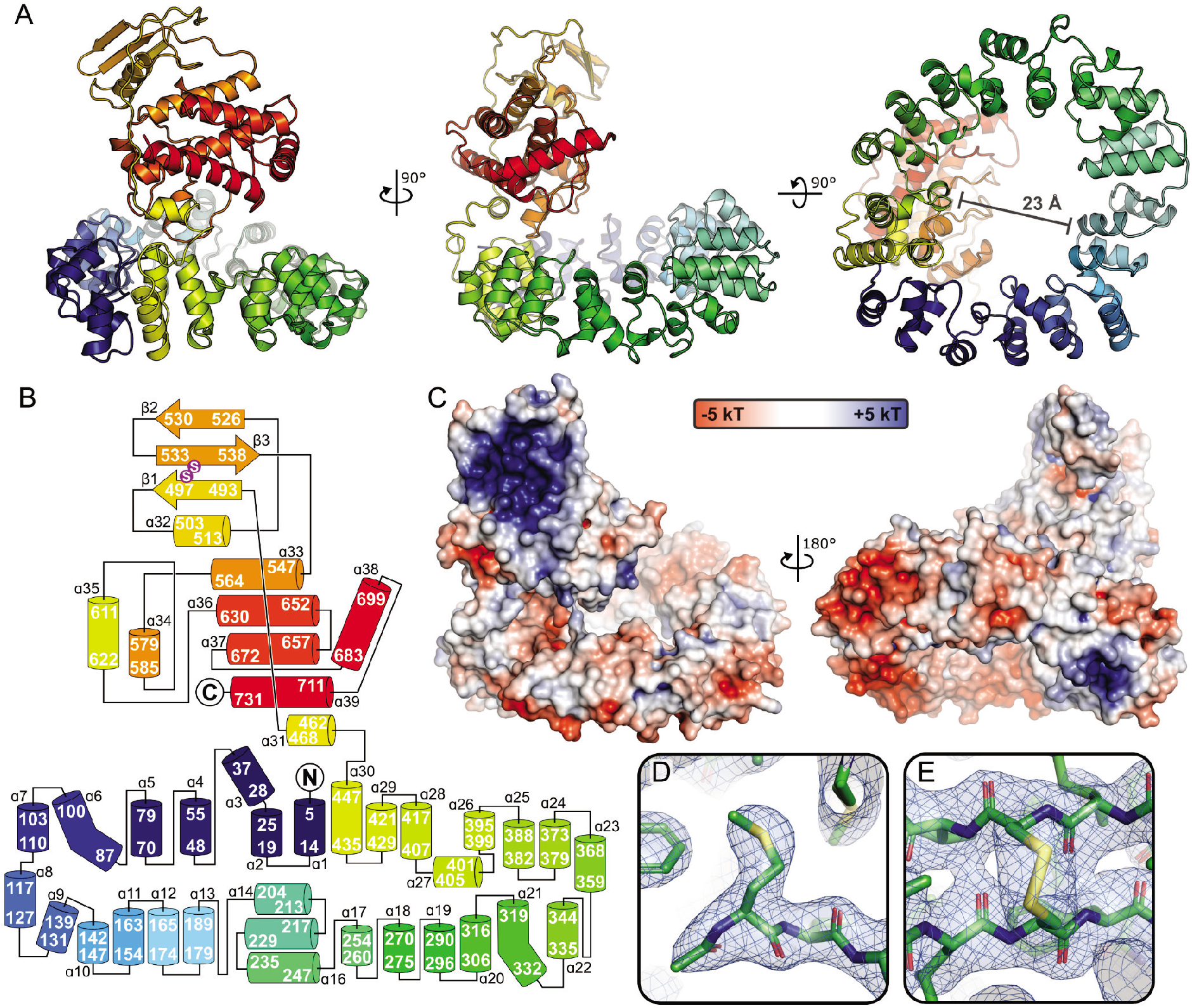
VACV E2 comprises novel N-terminal annular (ring) and C-terminal globular (head) domains. (A) The structure of VACV E2 is shown in three orthogonal views in ribbon representation, rainbow coloured from blue (N terminus) to red (C terminus). Molecular images were generated using PyMOL (Schrödinger LLC). The aperture of the ring domain is 23 Å wide at its narrowest point. (B) Schematic representation of VACV E2, with secondary structural elements coloured as in (A). Helices and sheets are shown as cylinders and arrows, respectively, with start and end residues shown. Sulphur residues that participate in a disulphide bond are shown in purple. (C) Molecular surface of E2 coloured by electrostatic potential from red (−5 kT) to blue (+5 kT), as calculated by APBS [52]. E2 is shown in two views, the left being rotated around the vertical and horizontal axes by approximately 15° from the middle panel of (A) to better illustrate the strong basic patch on the head domain and the lack of strong charge lining the centre of the ring domain. (D) The N-acetylated initiator methionine of E2 is shown in stick representation, with the final refined 2F_O_-F_C_ electron density map (1.2 σ) being shown as a blue semi-transparent mesh surface. (E) The disulphide bond between Cys residues 496 and 535 is shown in 2F_O_-F_C_ electron density (1.2 σ).

The N-terminal ring domain of VACV E2 is particularly striking, forming a central aperture that is 2.3 nm (23 Å) wide at its narrowest point (Figure 2A). While this is compatible with the diameter of B-DNA (2 nm), the radius is smaller than observed for DNA-binding proteins like PCNA and the inner surface of the ring domain lacks the positive electrostatic potential (Figure 2C) that would be expected for a DNA-binding protein [28]. The central aperture is too narrow to accommodate actin filaments (^~^6 nm diameter) or microtubules (^~^25 nm diameter), suggesting that the ring domain does not encircle cytoskeletal elements to promote VACV EV transport during infection. While VACV E2 is an acidic protein (theoretical isoelectric point 5.43 [29]) the head domain possesses a large basic patch on its surface (Figure 2C). Given the functional role of E2 in associating with intraluminal vesicles (ILVs) comprised of virions surrounded by trans-Golgi/early endosomal derived membranes, which are defined in part by their specific complement of phosphoinositides, it is tempting to speculate that the basic patch on the surface of the E2 head domain acts as a membrane recognition motif. Other noteworthy features of E2 include N-terminal acetylation of the initiator methionine (Figure 2D), which was verified by mass spectrometry, and the presence of a disulphide bond between Cys residues 496 and 535 in the head domain (Figure 2E) despite having included reducing agent (1 mM DTT) in the SEC purification buffer. Intramolecular disulphide bonding within cytosolic VACV proteins has been observed before [21] and its functional relevance remains unknown, although we note that a recent preprint has implicated redox proteins present in VACV lateral bodies in counteracting cellular oxidative stress generated during infection [30]. In summary, the structure of VACV E2 has provided some potential functional insights but, given the lack of structural homology to other well-characterised proteins, definitive mechanistic information has remained elusive.

Recently the application of deep learning technologies to the prediction of protein structures has made the resultant models significantly more accurate with regards to the backbone conformation (overall protein fold), although the side chain conformation prediction remains more challenging [31]. The lack of sequence identity between VACV E2 homologues and any other protein family, and the novelty of both the ring and head domains, makes E2 a particularly difficult target for structural prediction. Models of VACV E2 were thus generated using two leading structure prediction packages, AlphaFold2 (AF2) [32] and RoseTTAFold (RTF) [33]. Both AF2 and RTF accurately predicted the presence of two domains in the E2 structure, both successfully identifying that the head of E2 would have a compact globular fold while the ring would have an extended helical conformation.

Detailed analysis shows that predictions of the head domain by both AF2 and RTF were more accurate than for the ring domain (Figure 3A-D). Superpositions using SSM [25] and LGA (cutoff = 4 Å) [34] demonstrate that the head domain from the top ranked AF2 model can be superposed on the equivalent region of the experimental structure (residues 455–732, 278 residues in total) with a root-mean-squared deviation (rmsd) of 1.12 Å across 261 Cα atoms and a Global Distance Test Total Score (GDT_TS) of 87.2, which is remarkably accurate (Figure 3A). The largest discrepancy is at residues 477–492, which form part of the extended surface loop between the first helix and sheet of the head domain (Figure 3A). The prediction by RTF is less accurate (3.08 Å rmsd across 277 C^α^ atoms, GDT_TS = 49.8), consistent with accurate prediction of the gross topology but significant differences in the relative orientation of secondary structural elements (Figure 3B). With regards to the ring domain (residues 1–454), the top two AF2 models accurately predicted the ‘closed’ conformation of the E2 ring domain (Figure 3C) whereas the top RTF model predicted an ‘open’ conformation of the ring domain, the correct ‘closed’ conformation being observed in the second-ranked model (Figure 3D). The AF2 prediction was again more accurate (2.39 Å rmsd across 433 C^α^ atoms, GDT_TS = 68.3) than that of RTF (4.11 Å across 203 C^α^ atoms, GDT_TS = 36.7, for the top ranked model and 3.44 across 402 C^α^ atoms, GDT_TS = 45.3, for the second ranked model with a ‘closed’ ring). Furthermore, RTF demonstrated more heterogeneity in the conformation of the head domain relative to the ring. Neither AF2 nor RTF predicted the same relative domain orientation as seen in the experimental structure (Figure 3E), although it is possible that the two domains of E2 move relative to each other in solution and thus sample additional conformations not observed in the crystal structure.

**Figure 3.**
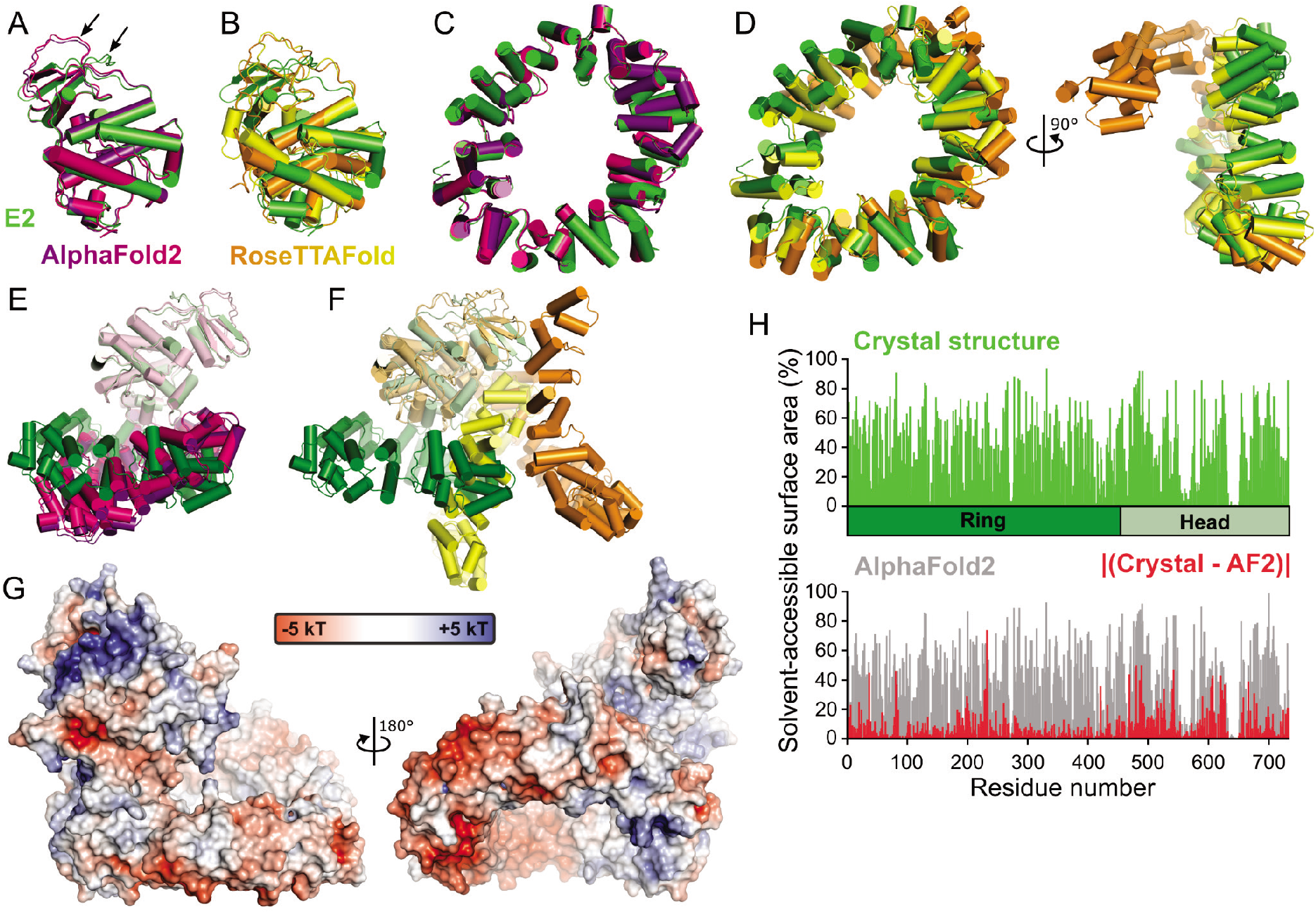
Assessment of prediction of the VACV E2 structure by AlphaFold2 (AF2) and RoseTTAFold (RTF). All superpositions were performed using SSM [25]. (A) Superposition of the head domain from the E2 crystal structure (green) with the top two models from AF2 (purple and pink, respectively). The loop between residues 477–492, where the backbone conformation of the models differs significantly from the crystal structure, is denoted with an arrow. (B) Superposition of the head domain from the E2 crystal structure (green) with the top two models from RTF (orange and yellow, respectively). (C) Superposition of the ring domain from the E2 crystal structure with the top two models from AF2, coloured as in (A). (D) Superposition of the ring domain from the E2 crystal structure with the top two models from RTF, shown in two orthogonal views and coloured as in (B). (E) and (F) Orientation of the E2 ring domain relative to the head domain in the top two AF2 (E; purple and pink) or RTF (F; orange and yellow) models compared with the E2 crystal structure (green). (G) Molecular surface of the top AF2 model of E2 coloured by electrostatic potential from red (−5 kT) to blue (+5 kT), as calculated by APBS [52]. E2 AF2 model is oriented as in Figure 2C. (H) Percent solvent accessibility of residues in the E2 crystal structure (green, top) or AF2 model (grey, bottom) as calculated using AREAIMOL [35, 36]. The absolute difference between calculated accessibility for the crystal structure and AF2 model is shown in red.

While the above analysis confirms that the AF2 model is closer to the crystal structure of E2 than that RTF model, the obvious use-case of *ab initio* modelling for molecular virologists is in situations where a reference crystal structure is not known. Such structural models can generate functional hypotheses by identifying structural homology to proteins of known function. They can also inform site-directed mutagenesis experiments by identifying surface-exposed residues and prominent surface features such as charged, hydrophobic or conserved patches. The question thus arises: *How useful is the AF2 model as a basis for generating hypotheses and designing mutations to test E2 function?* As mentioned above, the structures of the E2 ring and head domains do not share significant structural homology with other domains, but queries of the PDB with the two domains of the AF2 model using DALI [24] (after release of the PDB-deposited E2 structural coordinates) succeeded in identifying structural homology for each domain to the E2 crystal structure (Z = 26.5 and 31.1 for ring and head domains, respectively), suggesting that these domain models would have identified significant structural homologues should they have existed. The surface charge of the AF2 model is very similar to that of the crystal structure (compare Figure 3G with Figure 2C), although we note that the large basic patch on the head domain is less prominent due to the different conformation of residues 477–492 in the AF2 model. Furthermore, the percentage solvent-accessibility of each residue as calculated using the CCP4 program AREAIMOL [35, 36] was similar between the crystal structure and AF2 model (Figure 3E; Spearman nonparametric correlation coefficient ρ = 0.9253, P < 0.0001, as calculated using Prism7 [GraphPad]). Surprisingly, correlation was higher for the ring domain (ρ = 0.9475) than for the globular head domain (ρ = 0.8945), perhaps owing to its higher surface-area to volume ratio, but overall the AF2 model is clearly capable of predicting those residues that are buried in the core of the protein and should be avoided for mutagenic studies.

In addition to accelerating the generation and testing of functional hypotheses, high-quality structural models can simplify the process of solving macromolecular structures. Molecular replacement is a technique used for solution of the crystallographic phase problem whereby initial phases for a diffraction dataset are obtained from the atomic coordinates of a structure with a highly similar fold [37], rather than using heavy-atoms and/or anomalous scatterers to phase the structure as was done for E2. Molecular replacement represents a stringent test for the ‘value added’ by structural models [38], which has increased dramatically with the advent of deep learning techniques for protein structure prediction [31, 39, 40], and so the best-ranked models of E2 obtained from AF2 and RTF were tested for their ability to solve the structure of E2. The molecular replacement phasing experiment was performed using phenix.phaser [41] with a single search model (100% sequence identity to target structure), success being indicated by a translation function Z (TFZ) score > 8 [42]. The AF2 and RTF full-length models could not be used to solve the structure, nor could the ring domains of these models. The use of structural ensembles, where multiple models are superposed, can improve the signal in molecular replacement experiments by upweighting the phase contribution of structural regions confidently predicted to have the same conformation across multiple models (and are thus more likely to be correct) and downweighting the contribution from variable regions [41]. However, ensembles of the ring domains from the top 5 AF2 and RTF models generated by phenix.ensembler were also incapable of solving the structure. Similarly, the head domain of the top RTF model, or an ensemble of the head domains from the top 5 models, could not solve the structure. However, the head domain of the AF2 model could be successfully positioned in the crystallographic asymmetric unit (TFZ = 19.5) and using this model as a fixed component allowed positioning of the AF2 model ring domain (TFZ = 14.1). Subsequent automated structure completion using phenix.autobuild [43] confirmed that these molecular replacement solutions provided sufficient phase information for successful structure determination, the autobuild model comprising 737 residues with a *R*_free_ of 0.336 and rmsd of 0.81 Å across 697 Cα atoms when compared to the refined and deposited structure.

In conclusion, we have solved the crystal structure of VACV E2 to 2.3 Å resolution. E2 comprises two novel domains, an N-terminal annular (ring) domain and C-terminal head domain. While the fold of the head domain shares weak homology with cellular (pseudo)kinases and contains a large basic surface patch that may bind phosphoinositide headgroups, the lack of strong structural homology hampers attempts to infer E2 function by analogy to other proteins. Being a multi-domain protein with novel domain folds and limited availability of homologous protein sequences, VACV E2 is an excellent test for the ability of modern deep-learning algorithms to predict ‘difficult’ viral protein structures. The results are impressive, with both AF2 and RTF correctly predicting the overall fold of both domains and AF2 predicting the head domain with very high precision. AF2 models will prove a significant resource for the molecular virology community, allowing the identification of structural homologies in the absence of identifiable sequence homology, the exploration of protein surface features, and accelerating the experimental determination of novel viral protein structures.

## Acknowledgements

The authors thank Dr Len Packman (University of Cambridge) for performing mass spectrometry experiments, Mr Ben Butt (University of Cambridge) for assistance with installing AlphaFold2, SBGrid for performing the Wide-Search Molecular Replacement experiment, and Tristan Croll (University of Cambridge) for reading the draft manuscript. We thank Diamond Light Source for access to beamlines I04-1 (mx11235) and I04 (mx15916), remote access to which was supported in part by the EU FP7 infrastructure grant BIOSTRUCT-X (Contract No. 283570). This work was supported by a Wellcome Trust Principal Research Fellowship (090315) to GLS, an MRC research grant (MR/R010536/1) to GLS and DC, a Royal Society University Research Fellowship (UF100371) to JED and a Sir Henry Dale Fellowship (098406/Z/12/B), jointly funded by the Wellcome Trust and the Royal Society, to SCG.

## Notes

http://doi.org/10.2210/pdb7PHY/pdb

https://doi.org/10.17863/CAM.74391

### Competing Interest Statement

The authors have declared no competing interest.

